# TAMIPAMI: Software and methods for PAM/TAM identification in CRISPR and OMEGA gene editing systems

**DOI:** 10.64898/2026.05.15.725432

**Authors:** Carlos Orosco, Piyush K. Jain, Adam R. Rivers

## Abstract

Protospacer adjacent motifs (PAMs) and target-adjacent motifs (TAMs) are essential for target recognition by CRISPR-Cas and TnpB nucleases. Here we present TAMIPAMI, an efficient experimental and computational framework for rapid PAM/TAM identification. TAMIPAMI requires only a single control library and Cas or TnpB-treated library, simplifying experimental design, reducing cost, and providing greater accessibility for users. The platform interprets sequencing data with interactive visualizations and introduces a novel algorithm that determines the minimal exact set of degenerate IUPAC sequences describing the observed PAM/TAM patterns. Using this approach, we accurately recovered canonical motifs for several nucleases, including SpCas9, LbCas12a, AsCas12a, BrCas12b, Cas12i1, and AmaTnpB. TAMIPAMI is available as both a web application and command-line tool, ultimately providing an accessible and efficient platform for PAM/TAM discovery and characterization across CRISPR and OMEGA systems.

## Main

CRISPR-Cas systems have revolutionized genome editing and diagnostics by enabling precise, RNA-guided nucleic acid cleavage. In nature, CRISPR-associated (Cas) proteins use CRISPR RNAs (crRNAs) to recognize and cleave foreign DNA or RNA, a mechanism now widely repurposed for genome engineering and molecular diagnostics.^1,2^ These systems are highly diverse, spanning multi-effector Class 1 complexes, single-effector Class 2 nucleases, and related RNA-guided enzymes such as OMEGA-family TnpB, which are evolutionarily linked to Type V (Cas12) systems.^3–5^ TnpBs are compact nucleases that utilize a longer ωRNA, making them attractive platforms for engineering.

A key determinant of targeting is the requirement for a short sequence motif adjacent to the target: the protospacer adjacent motif (PAM) in CRISPR nucleases or the target-adjacent motif (TAM) in TnpBs. These motifs are essential for target recognition and activation and ultimately define the sequences accessible to each nuclease. Accurate identification of PAM/TAM requirements is therefore critical for characterizing activity and expanding targeting scope.

A range of computational and experimental approaches have been developed for PAM/TAM identification. In silico methods analyze genomic datasets and CRISPR spacer alignments to predict candidate motifs, enabling rapid initial screening.^6,7^ Experimental approaches utilizing bacterial depletion assays, synthetic circuits, and in vitro transcription–translation (TXTL) systems, provide functional validation and enable high-throughput characterization under controlled conditions.^8^ Mammalian cell-based assays further define nuclease activity in vivo through sequencing or reporter-based readouts. ^9–15^

Among these, HT-PAMDA^16^, has emerged as a particularly powerful and comprehensive approach. This method directly measures nuclease cleavage across large, randomized PAM libraries using massively parallel sequencing, enabling quantitative and high-resolution mapping of PAM preferences. By capturing cleavage outcomes across thousands of sequences simultaneously, HT-PAMDA provides a detailed view of nuclease specificity and has become a valuable framework for PAM discovery. However, current implementations require multiple time-point libraries and rely on monolithic analysis pipelines with rigid assumptions about library design and sequencing structure, limiting accessibility, and broader adoption.

To address these limitations, we developed TAMIPAMI, a flexible and accessible framework for PAM/TAM identification. TAMIPAMI provides key analytical improvements for high-throughput nuclease cleavage assays. Standard methods rely on normalized k-mer counts to estimate depletion relative to a control library, but sequencing data are inherently compositional, meaning each k-mer count is interdependent^17^. To address this, TAMIPAMI applies a centered log-ratio (CLR) transformation, which reduces variance among non-cleaved k-mers in noisy datasets and improves detection of true PAM/TAM signals.

TAMIPAMI also introduces a novel algorithm to compute the exact minimal set of degenerate sequences describing nuclease specificity. Given observed PAM sequences, the method identifies the smallest collection of IUPAC-degenerate codes that completely and exactly covers them, rather than approximating a consensus motif. This formulation corresponds to a variant of the NP-hard Exact Cover Set Problem, but with important extensions: candidate solutions are degenerate nucleotide patterns rather than discrete subsets, each pattern implicitly representing many sequences. As a result, the search space grows combinatorially (15^k for length k), while also being constrained by feasibility (patterns must be derivable from observed sequences) and redundancy (more general patterns can subsume specific ones). These competing constraints make direct application of classical exact cover solvers insufficient and require tailored strategies for pruning and representation.

To solve this, TAMIPAMI uses a two-phase approach. First, it groups similar sequences and generates candidate degenerate patterns through local expansion, maximizing coverage while avoiding false positives. Second, it applies global optimization to select the minimal set of patterns that exactly covers all observed sequences.

Overall, TAMIPAMI is a user-friendly and automatable framework for analyzing nuclease cleavage data, combining rigorous statistical analysis with a principled approach for representing complex PAM/TAM landscapes. Available as both a web application and command-line tool, it requires only a single control and experimental library, reducing experimental complexity and cost. Using this approach, we accurately characterized PAMs for SpCas9, LbCas12a, AsCas12a, BrCas12b, and Cas12i1, as well as the TAM for AmaTnpB, with laboratory data, enabling improved discovery, engineering, and characterization of CRISPR and OMEGA nucleases.

## Results

### Development of a streamlined PAM/TAM screening library

To generate input data for the TAMIPAMI algorithm, we developed a streamlined screening method based on the protocols described by Walton et al. and Karvelis et al.^8,16^ Our goal was to establish a simplified approach in which a single dataset is sufficient for both PAM and TAM identification (Fig. 1a). We began by using the PAM plasmid libraries from Walton et al., in which the target site is flanked by eight randomized nucleotides (8×N) positioned either at the 3′ or 5′ end. The location of these randomized nucleotides is critical, as Type II (Cas9) systems recognize PAM sequences at the 3′ end of the target site, whereas Type V (Cas12) and TnpB systems require PAM/TAM recognition at the 5′ end.

**Figure 1.**
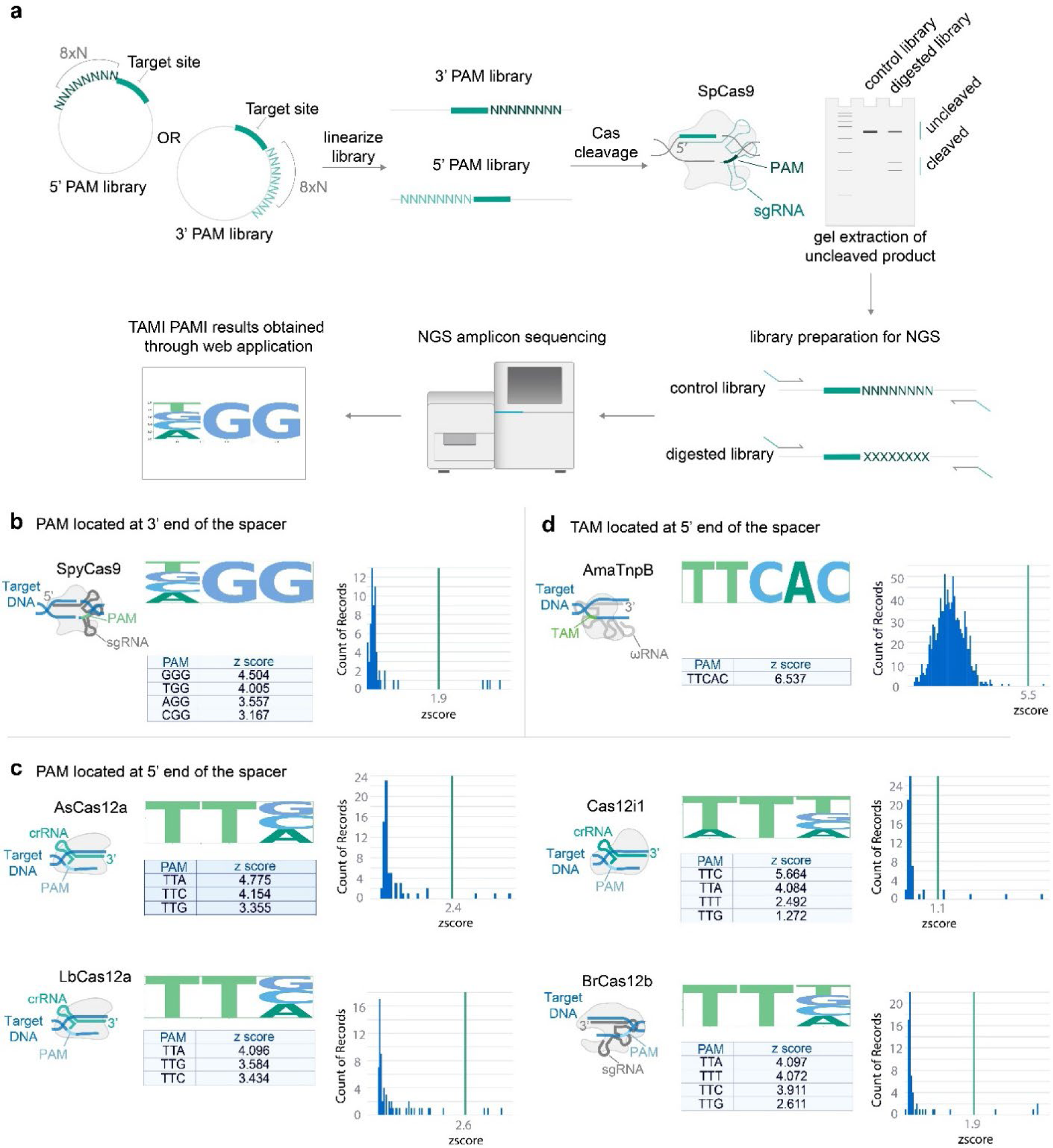
PAM and TAM motifs identified by TAMIPAMI for six nucleases. **(a)** Schematic overview of the TAMIPAMI experimental workflow for PAM/TAM identification. Plasmid libraries containing randomized PAM or TAM regions (8xN adjacent to the target site—at the 3′ end for Cas9 or at the 5′ end for Cas12 and TnpB) were acquired from Addgene and linearized using the PvuI-HF restriction enzyme. In vitro cleavage assays for Cas proteins were performed at a 10:10:1 molar ratio of Cas protein, crRNA, and linearized target DNA. For AmaTnpB the ratios were 60:60:1. After the reaction, uncleaved plasmid fragments were separated on a 1% agarose gel. Both the uncleaved bands and untreated control libraries were gel-extracted and PCR-amplified to add Illumina adaptors and barcodes. Paired-end sequencing (2 × 150 bp) was performed on an Illumina MiSeqDx. FASTQ files for both uncleaved and untreated control libraries were uploaded to the TAMIPAMI web application. **(b)** 3-mer motif logos for SpCas9 showing PAM sequences at the 3′ end of the target site. **(c)** 3-mer motif logos for Cas12 family nucleases (LbCas12a, AsCas12a, BrCas12b, Cas12i1) showing PAM sequences at the 5′ end. **(d)** 5-mer motif logo for AmaTnpB showing the TAM sequence at the 5′ end of the target site. The motifs were identified using the TAMIPAMI web application. Libraries containing randomized 8xN sequences adjacent to the target site were processed following the workflow described in

**Figure 2.**
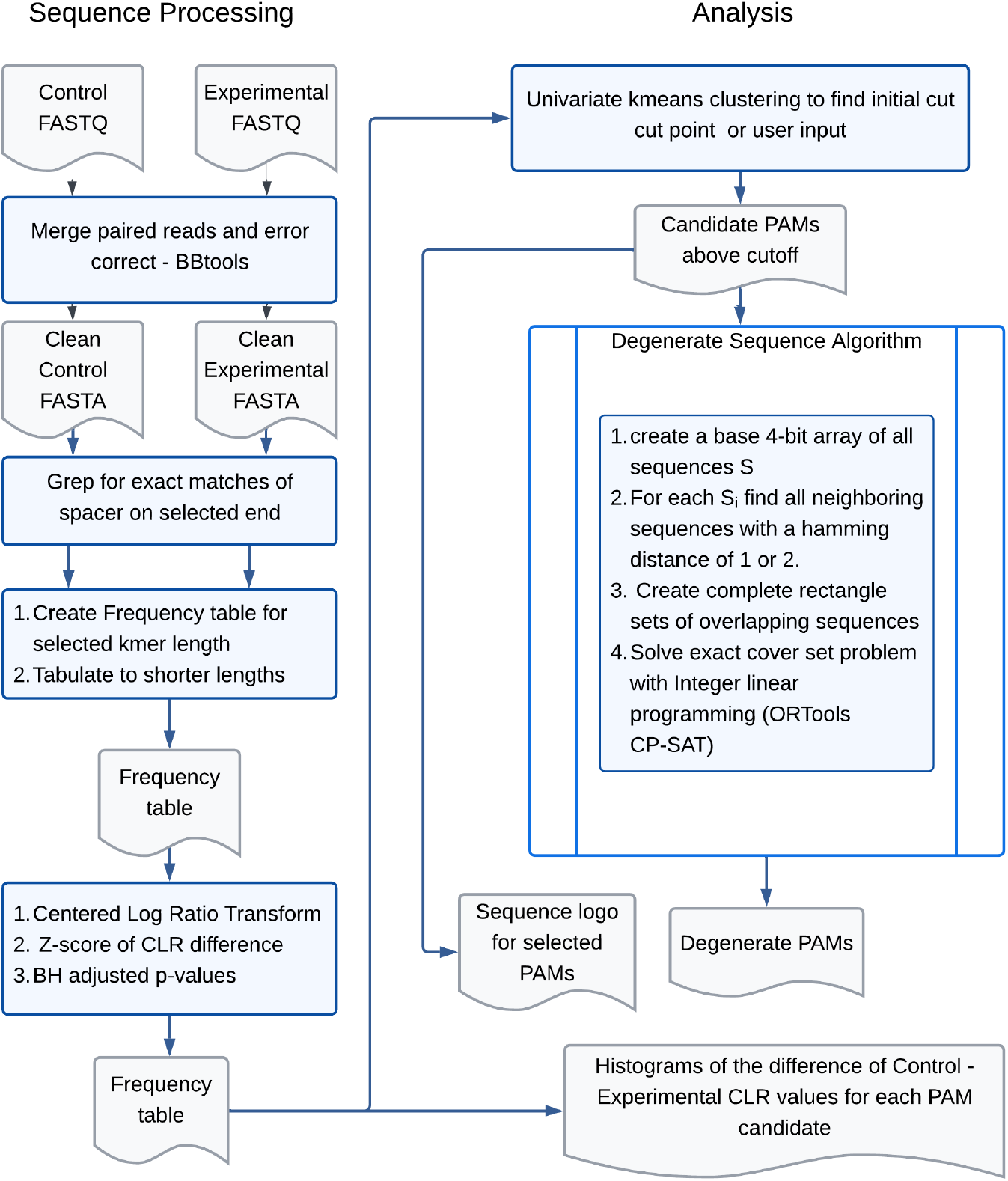
TAMIPAMI algorithm flowchart. In the processing phase, paired end reads are merged for error correction; whole single-end reads are processed directly. Exact spacer matches are found, and the adjacent sequence is counted for a control and a digested sequence library. A frequency table is constructed then grouped to calculate the frequencies for putative PAMs/TAMs of shorter lengths down to 3nt. The data are CLR transformed, and a z-score of the CLR difference is calculated along with BH corrected P-values for that sequence relative to the population. In the analysis phase, an initial split between the cut and uncut PAM/TAM sequences is suggested using a univariate k-means algorithm. The user can inspect and refine this cut point. All sequences above this cut point are then provided to the degenerate sequence algorithm which computes a minimal exact IUPAC degenerate DNA sequence set describing the PAM/TAM sequences cleaved.

After obtaining these plasmids, we linearized them using the restriction enzyme PvuI to facilitate downstream processing. The linearized libraries were then subjected to digestion with either Cas or TnpB nucleases, pre-complexed with their respective guide RNAs. This allowed selective cleavage at sites containing recognized PAM/TAM sequences. Both digested and undigested control libraries were run on agarose gels, and the uncleaved bands were gel-extracted from each condition for further processing. The extracted DNA was then amplified in two successive PCR steps to add sequencing adaptors and barcodes compatible with Illumina platforms.

### TAMIPAMI experimental validation with Cas and TnpB

The TAMIPAMI algorithm was validated using experimental data from SpCas9 (Fig. 1b), LbCas12a, AsCas12a, BrCas12b, Cas12i1 (Fig. 1c), and AmaTnpB (Fig. 1d). Each enzyme was tested using the protocol described above, and the resulting sequencing files were uploaded to the TAMIPAMI web application. In all cases, the algorithm accurately identified the canonical PAM or TAM sequences reported in the literature, achieving 100% concordance. SpCas9 cleavage was replicated to confirm the reproducibility of the results.

After uploading data to the web interface, the application processes the reads, merging paired end reads and counting the number of PAM/TAM sites between three and the maximum length provided by the user. The data are CLR transformed, and the z-score of the Experimental – Control CLR data are displayed as a histogram. Uncleaved sequences form a cluster near 0, while cleaved sequences have higher z-scores. At each length, TAMIPAMI uses 1-dimensional k-means clustering to propose a separation for cleaved sequences. The user can adjust the separation value interactively. After each adjustment, the app creates a new sequence motif logo, a new set of degenerate sequences, and a filtered table of results. The user can also switch between tabs with different kmer lengths to determine the optimal length of the PAM/TAM site. Figure 1 shows the results of TAMIPAMI and the true PAM/TAM sequences.

In addition to motif visualization via sequence logos, TAMIPAMI provides the z-score for each individual PAM or TAM, offering a quantitative measure of the depletion strength. For instance, in the case of SpCas9, the canonical NGG PAM was identified, with the motif logo and z-score analysis revealing that GGG exhibits the strongest cleavage preference among the NGG variants. Higher z-scores indicate strong depletion and thus specific cleavage activity at those motifs.

Notably, TAMIPAMI achieves these results using only a single pair of experimental and control datasets, without requiring time-course sampling or biological replicates. This streamlines the experimental workflow while still delivering accurate and statistically robust motif identification.

### Performance benchmarking

Both the command line application and the web application versions of TAMIPAMI were benchmarked for run time, memory usage and CPU usage on a 2023 MacBook Pro M2. The results are in Table 1 and Table 2. Samples with between 302,816 and 2,965,144 read pairs could be processed in 11-34 seconds. The process step averaged 5-11% CPU usage and 262-268MB of memory usage, while predicting step averaged between 80-157% of CPU and 319-350MB of memory usage. Web application times were similar, ranging from 4-25 seconds with negligible client memory and CPU usage.

**Table 1.** Command Line Benchmark Results.

| Command Line Benchmark Results |  |  |  |  |  |  |  |  |  |
| --- | --- | --- | --- | --- | --- | --- | --- | --- | --- |
| Experiment | Enzyme | Process Time (sec.) | Predict Time (sec.) | Process Max Mem (MB) | Predict Max Mem (MB) | Process Avg CPU (%) | Predict Avg CPU (%) | Control Read pairs | Experimental Read pairs |
| 1 | SpyCas9 | 14.62 ± 0.22 | 4.82 ± 0.19 | 268.28 ± 1.07 | 317.17 ± 4.47 | 11.56 ± 0.28 | 89.98 ± 2.67 | 642,368 | 551,539 |
| 2 | SpyCas9 | 18.39 ± 0.33 | 4.26 ± 0.15 | 268.50 ± 4.34 | 349.84 ± 3.78 | 9.55 ± 0.56 | 157.23 ± 14.28 | 910,598 | 827,259 |
| 3 | AmaTnpB | 7.56 ± 0.91 | 3.78 ± 3.11 | 262.31 ± 5.77 | 319.69 ± 3.60 | 23.87 ± 2.22 | 80.06 ± 16.73 | 171,537 | 130,279 |
| 4 | AsCas12a | 27.42 ± 0.32 | 4.13 ± 0.13 | 268.49 ± 4.88 | 319.22 ± 3.57 | 6.31 ± 0.23 | 88.76 ± 3.02 | 642,982 | 2,247,820 |
| 5 | BrCas12b | 14.79 ± 0.20 | 7.54 ± 3.48 | 267.12 ± 1.43 | 315.75 ± 3.32 | 11.13 ± 0.27 | 88.09 ± 14.51 | 642,982 | 686,819 |
| 6 | Cas12i1 | 29.11 ± 0.18 | 4.56 ± 0.06 | 268.37 ± 2.46 | 317.43 ± 2.65 | 5.86 ± 0.26 | 91.74 ± 0.94 | 1,278,593 | 1,686,551 |
| 7 | LbCas12a | 20.98 ± 0.24 | 4.63 ± 0.17 | 268.47 ± 1.12 | 319.63 ± 3.70 | 8.05 ± 0.22 | 89.29 ± 3.81 | 642,982 | 1,436,737 |

**Table 2.** Chrome App Benchmark Results.

| Chrome App Benchmark Results |  |  |  |  |  |  |
| --- | --- | --- | --- | --- | --- | --- |
| Experiment | Enzyme | Elapsed Time (sec.) | Max Mem (MB) | Avg CPU (%) | Control Read pairs | Experimental Read pairs |
| 1 | SpyCas9 | 12.10 ± 0.13 | 14.82 ± 0.33 | 0.0000 ± 0.0000 | 642,368 | 551,539 |
| 2 | SpyCas9 | 15.97 ± 0.26 | 14.61 ± 0.06 | 0.0000 ± 0.0000 | 910,598 | 827,259 |
| 3 | AmaTnpB | 4.00 ± 0.12 | 16.17 ± 0.90 | 0.0006 ± 0.0019 | 171,537 | 130,279 |
| 4 | AsCas12a | 25.54 ± 0.53 | 14.59 ± 0.02 | 0.0000 ± 0.0000 | 642,982 | 2,247,820 |
| 5 | BrCas12b | 12.36 ± 0.11 | 14.61 ± 0.04 | 0.0000 ± 0.0000 | 642,982 | 686,819 |
| 6 | Cas12i1 | 27.37 ± 0.17 | 11.89 ± 2.37 | 0.0000 ± 0.0000 | 1,278,593 | 1,686,551 |
| 7 | LbCas12a | 18.42 ± 0.16 | 10.01 ± 0.01 | 0.0000 ± 0.0000 | 642,982 | 1,436,737 |

### Comparison to HT-PAMDA

TAMIPAMI and HT-PAMDA operate somewhat differently, but we compared the results of a dataset provided by the HT-PAMDA paper to the results of that data processed in TAMIPAMI. HT-PAMDA uses one undigested control library and three digested libraries at different timepoints. For the comparison, we selected the control library and the last timepoint of the experimental library to process with TAMIPAMI. The count of 4mers adjacent to the spacer for their control library was identical in HT-PAMDA in TAMIPAMI. In their experimental library A length definition error on line 288 of the file PAMDA.py caused their experimental data counts report a mean of 35.6% ± 16.0 S.D. of the total reads present. We were able to analyze the forward control and last timepoint reads of their experimental data with TAMIPAMI. Their forwards reads were 65 pb and their reverse reads were 10pb so error correction by merging could not be done by TAMIPAMI as it can when forward and reverse reads are the same length. Histograms of their data indicated an enrichment of DRR ({G, T, A}, {G, A}, {G, A}}) PAMs which is similar to NGG. but there was not a clear bimodal distribution in their data when processed through TAMIPAMI (Fig. S1). HT-PAMDA and TAMIPAMI process data differently, which is discussed below.

## Discussion

In this study, we present TAMIPAMI, a user-friendly computational tool designed to accurately identify PAM and TAM sequences for CRISPR-Cas and TnpB systems. Building on the framework established by Walton et al. in their HT-PAMDA protocol, we sought to streamline both the experimental and computational components to reduce complexity and data requirements. Unlike existing approaches, TAMIPAMI eliminates the need for time-course sampling or biological replicates, relying solely on a single pair of digested and control libraries. All laboratory steps are performed in vitro, significantly simplifying the workflow. A key strength of TAMIPAMI lies in its minimal input requirements and accessibility. The entire pipeline is integrated into a web-based interface, removing the need for package installations or coding experience. This lowers the barrier for widespread adoption and enables rapid motif identification in diverse systems. TAMIPAMI is also available as a command-line application for users wanting to integrate the software into existing bioinformatic workflows.

In our comparison, TAMIPAMI and HT-PAMDA differ somewhat in their results. While TAMIPAMI clearly shows it can distinguish the NGG PAM in our data, it found a less specific DRR motif when analyzing time point 3 of the HT-PAMDA SpCas9 data. This may have been due to incomplete cleavage of that library. HT-PAMDA does not output a simple histogram of sequences, but it does output heatmaps of rates that appear consistent with a DRR or NGG motif. TAMIPAMI uses a simple model to detect cleavage which accounts for the compositional nature of the data and makes only the weak assumption that the PAM/TAM targets a minority of the possible kmers. HT-PAMDA quantifies kmer depletion by comparing an uncleaved control library (representing Time 0) against experimental samples collected at 1, 8, and 32 minutes. For each timepoint, the pipeline first calculates the relative abundance of every kmer to account for sequencing depth. To correct for the compositional “inflation” of inactive sequences, it identifies the five kmers with the most positive linear slopes across all timepoints, using their median as a scaling factor to ‘detrend’ the entire dataset. Finally, the pipeline applies a two-parameter exponential decay function to each kmer, using ordinary least squares to derive a cleavage rate constant k. There are a few methodological complications with this approach.

In revisiting these data, we noted an error in counting the timepoint reads reduced those counts by about 65%, making it appear as if there was a rapid cleavage of all kmers followed by slower reductions. The second step was to fit an ordinary least squares linear regression to the timepoints and identify the five kmers with the largest slopes (least cleavage) then use that to normalize all other slopes. This normalization attempts to de-trend the universal downward trend caused by the counting issue. For NGG, 60 of 64 3-mers are expected not to be cleaved, but only the 5 least-cleaved were selected for normalization, potentially introducing additional variability into the data. This de-trended data is then used to fit an ordinary least squares exponential decay equation. In those fits, the y-intercept parameter a was allowed to vary, although in principle it should be fixed to 1 (the normalized control library abundance) there are only 4 points and 2 degrees of freedom k *and* a can be changed to fit the data, fixing a would reduce noise and enforce the biological reality of a 100% starting concentration. The use of ordinary least squares rather than weighted least squares fails to add additional credence to higher count samples, but in this case the libraries had similar counts, so that effect is less pronounced in this data. The general trend of the cleavage assay still comes through with the approach used by HT-PAMDA, likely due to the strength of the biological signal. TAMIPAMI, in contrast, has a simpler endpoint focused approach, which clearly distinguishes the PAM/TAM in 6 test datasets. We chose the endpoint approach because for routine screening we are most interested in CRISPR enzymes which clearly show strong cleavage under normal experimental reaction times.

Importantly, TAMIPAMI identifies canonical PAM/TAM motifs and provides quantitative rankings of sequence efficiency based on z-score. For example, in the case of SpCas9, TAMIPAMI confirmed the known NGG consensus, and further revealed GGG as the most efficiently cleaved PAM which is consistent with prior experimental studies. This demonstrates that TAMIPAMI can recover quantitative distinctions in PAM efficiency without the need for additional optimization or protocol changes, providing users with interpretable, statistically robust outputs.

Another valuable feature of the platform is its novel, degenerate sequence resolution tool, which identifies the minimal degenerate motif that accurately represents all high-confidence hits. This functionality reduces ambiguity and helps researchers pinpoint comprehensive PAM or TAM sets in a clear, compact format.

TAMIPAMI also expands beyond well-established Cas systems to include novel enzymes such as TnpBs, which are known to have more stringent TAM requirements. For instance, AmaTnpB, shown in Figure 3, requires a five-nucleotide TAM that was accurately captured by our algorithm. This highlights the algorithm’s ability to resolve complex motif architectures, even for compact nucleases with less-characterized PAM/TAM sequences.

Ultimately, our goal with TAMIPAMI was to create an end-to-end solution from streamlined experimental design to a robust and intuitive analysis platform for fast, accurate, and accessible PAM/TAM identification. By offering high resolution, reduced complexity, and platform independence, in a way that is complementary to existing time-resolved methods such as HT-PAMDA, TAMIPAMI serves as a powerful resource for the continued discovery, characterization, and engineering of CRISPR-Cas and TnpB systems.

## Methods

### Protein expression and purification

For proteins AsCas12a, LbCas12a, BrCas12b and Cas12i1, the following protocol was used for protein expression and purification. SpyCas9 was purchased from New England Biolabs (#M0646T).

Rosetta cells were transformed with the desired plasmid and cultured overnight at 37°C on agar plates supplemented with Ampicillin or Kanamycin. One colony was later expanded on 10 ml of LB medium for 12 hours. The culture was later added to 2 L of TB medium and grown to an optical density of 0.6-0.8. Once OD was reached, the culture was cooled 30 minutes on ice and then IPTG was added to a final concentration of 0.5 mM. Finally, the culture was incubated 18 hours at 16C.

After incubation, cells were centrifuged at 10,000 xg for 10 minutes. The media was discarded, and the pellet was resuspended in Lysis Buffer (500 mM NaCl, 50 mM Tris-HCl, pH 7.5, 20 mM Imidazole, 0.5 mM TCEP, 1 mM PMSF, 0.25 mg/mL Lysozyme, DNase I). The mixture was later sonicated for 30 minutes and then centrifuged at 40,000 xg for 30 minutes. The supernatant was collected and filtered through a 0.22 µm syringe filter (Cytiva, Catalog #9913-2504). The lysate was then purified through a BioLogic DuoFlow™ FPLC system (Bio-Rad) with a 5 ml Histrap FF column (Cytiva, Catalog #17525501, with Ni2+ replaced by Co2+). The proteins were eluted in Buffer B (500 mM NaCl, 50 mM Tris-HCl, pH 7.5, 250 mM Imidazole, 0.5 mM TCEP).

For all proteins, except Cas12i1, the eluted fractions were merged and dialyzed in a 10 kDa – 14 kDa MWCO bag with TEV protease (sourced from David Waugh, Addgene #8827). The dialysis bag was immersed in Dialysis Buffer (500 mM NaCl, 50 mM HEPES, pH 7, 5 mM MgCl2, 2 mM DTT) and stirred at 4°C overnight. Then the proteins were concentrated to approximately 2 mL using a 30 kDa MWCO Vivaspin® 20 concentrator. This concentrate was then balanced with 10 mL of Buffer C (150 mM NaCl, 50 mM HEPES, pH 7, 0.5 mM TCEP). Then the proteins were further purified through FPLC with a 5 mL Hitrap Heparin HP column. A gradient flow alternating between Buffer C and Elution Buffer D (2000 mM NaCl, 50 mM HEPES, pH 7, 0.5 mM TCEP) facilitated protein elution. Finally, buffer was exchanged in a 30 kDa MWCO Vivaspin® 20, transferred to a Storage Buffer (25mM Tris-HCl, 150mM NaCl, 2mM TCEP @ pH 8) then proteins were frozen at -80C for future use.

AmaTnpB was purified as described in Altae-Tran et al.

### In vitro cleavage assay

The in vitro cleavage assay is a simplified version of Russell. et al. protocol. In short, PAM libraries with 8xN nucleotides adjacent to the target site were acquired from Addgene (RTW554, RTW555, RTW572, and RTW574). Libraries RTW554, RTW555 have the PAM sequence at the 3’ end of the target site. These were used for Cas9 cleavage. Libraries RTW572, RTW574 have the PAM at the 5’ end of the target and used for Cas12 and TnpB cleavage. To start, 10 ug of plasmid libraries were linearized with 40 U of PvuI-HF restriction enzyme in NEB CutSmart buffer in a 40 ul reaction. The samples were incubated for 4 hours at 37C to achieve full digestion. The product was then purified with Beckman Coulter SPRI magnetic beads. 1.5x volume of beads were added to the linearized libraries, incubated for 5 minutes and placed in a magnetic rack. All liquid was removed and then beads were washed twice with 200 ul of 70% (v/v) ethanol. After ethanol removal, beads were left to dry until no ethanol was present. Finally, samples in the beads were eluted with 40 ul of nuclease-free water.

Library depletion was performed at 10:10:1 molar ratio of Cas protein, crRNA, and target. 40 ul reactions were prepared with 80 nM Cas9 or Cas12 and 80 nM crRNA reaction buffer (NEB Buffer 3.1 for SpCas9 or NEB Buffer 2.1 for all Cas12). The protein and crRNA were incubated at room temperature for 10 min. and then the linearized library was added to a final concentration of 8 nM. Samples were incubated at 37C for SpCas9, Cas12i1, LbCas12a, AsCas12a and 60°C for BrCas12b.

After incubation, the Cas-depleted libraries and a sample of the untreated library were loaded into a 1% agarose gel. The bands at the library size (~2.9 kb) were gel extracted with NEB Gel Purification Kit to posteriorly use for NGS sequencing.

### NGS library preparation for DNA sequencing

To sequence the untreated and depleted libraries, the gel purified products were PCR amplified to add adaptors and barcodes for Illumina NGS sequencing. This method again follows Walton. et al., PCR protocol with modifications to the primer sequences. In brief, the purified libraries were added to 25 ul Q5 polymerase PCR reaction to a final concentration of 300 pM. The Q5 reaction mixture contained the following, 500nM of PCR1 primers (each), 200 uM dNTPs, 0.5 M Betaine, 3 mM MgCl2, and 0.5 U of Q5 polymerase in Q5 reaction buffer. Samples were thermocycled for 30 cycles and then purified with SPRI beads as detailed previously. Purified products were again amplified with barcoded primers for a second round of PCR. For this step, 0.250 ng of the product was added to a standard Q5 PCR reaction as per manufacturers protocol with PCR2 primers. The samples were thermocycled for 10 cycles only. Lastly, the final library of amplicons was purified once again with SPRI beads and then pooled together at equal amounts and loaded into an Illumina MiSeqDx with a MiSeq Micro Kit v2 300 cycles (Illumina #MS-102-2002). MiSeq was set to perform 150×2 pair-end reads. FASTQ files were then uploaded to the web application for PAM identification analysis.

### Computational methods

TAMIPAMI makes three assumptions that allow it to detect sequences with only one control and one experimental library. The first assumption is that the endonuclease being characterized recognizes a small number of the total possible TAM/PAM sites. For example, the canonical SpCas9 PAM ‘NGG” would match 1/16 of all random 3-mers. This means that the distribution of kmer abundances is bimodal, and the mean is closest to the uncut population. The second assumption is that a useful guided endonuclease elicits strong, specific nuclease activity. This reduces the target sequence in the sequence library well above the background distribution of other kmer abundances. Finally, it is assumed that the random region of the plasmid library is reasonably random. These assumptions allow us to take advantage of the replication in the sequence population to statistically infer the PAM/TAM sequence the endonuclease recognizes.

TAMIPAMI initially processes sequence data into count tables. It takes, optionally gzipped, paired-end or single-end FASTQ sequence files for the control (undigested) and experimental (endonuclease-treated) library. The paired-end reads are merged and error-corrected using BBtools BBmerge.^18^ Reads are read with Biopython^19^ and exact matches to the spacer are identified with the Python regular expressions library. The user provides a maximum sequence length for the PAM/TAM (6 by default). TAMIPAMI generates a list of all possible canonical and non-canonical kmers at this length, then counts the number of times they appear in the control and experimental sample. These data counts are then merged to shorter length kmers down to length 3 by dropping the position distal to the spacer (the leftmost base for 3-prime orientation and the rightmost base for 5-prime orientation). This requires a single pass through the sequence data. The counts from the sequencing experiments are compositional count data^17,20^ and exist in the simplex number space. Practically, this means that the count of each kmer affects the counts of all other kmers because each kmer consumes the fixed number of reads available in a sequencing run. TAMIPAMI applies a CLR transformation to the simplex data, converting it to the real number space where z-scoring and other standard statistical methods can be applied.

Because this step only needs to be completed once, the processing step is a standalone subcommand in the command line version, and results are cached in the web application. The second step in identifying the PAM/TAM site is to select an appropriate cutoff value for the sequences that divides PAM/TAM sites from non-recognized sites. By default, TAMIPAMI performs optimal 1-Dimensional k-means clustering using the method of Song et al. 2011 implemented in the Python package ckmeans^21^. This generally provides a good starting point for separating the sites cleaved by the nuclease, but the web application provides a slider that can be used to modify the cut point. The web application also provides an interactive histogram, a table of filtered results, a sequence logo^22^ created with the Python Logomaker package, and a list of minimal degenerate sequences describing the set of PAM/TAMs above a cutoff value.

One key feature of TAMIPAMI is its ability to generate the minimal exact set of degenerate IUPAC sequences from a set of input TAM/PAM sequences that are cleaved by the test enzyme. TAMIPAMI uses a two-step approach to identify the degenerate sequence set. First, candidate degenerate sequences with exact coverage are created by greedy search of similar sequences, then the OR-Tools CP-SAT solver applies constraint programing optimization to merge those groups into a minimal exact degenerate set.

For candidate generation, the set is first encoded as a direct-address table indexed by base-4 encoded kmers, enabling constant-time membership checks. Then, seed patterns are generated using the Hamming neighborhood of ≤ 2 for each sequence. For each sequence, all of its neighbors are found, and each neighbor forms its own minimal seed pattern with the query sequence. For example, in the set {AAA, AAT, AAG, CCC}, we start with AAA as the query sequence. AAT and AAG have a Hamming distance to AAA of 1. So, we consider two pairs (AAA, AAT) and (AAA, AAG), which can be represented as {{A}, {A}, {A, T}} and {{A}, {A}, {A} {A, G}}. Each seed pattern is then independently expanded using greedy closure, which tests every possible addition of bases at every position and accepts an addition only if all sequences implied by the expanded pattern, not just those in the Hamming neighborhood, are present in the dataset. The process iterates until no further bases can be added without introducing sequences that were not observed, resulting in a *maximal* exact degenerate pattern.

The final selection of degenerate patterns is performed using the OR-Tools CP-SAT solver, an exact anytime integer optimizer. CP-SAT introduces one Boolean decision variable per candidate pattern and enforces exact-cover constraints requiring every input sequence to be covered by exactly one selected pattern. The solver then minimizes a hierarchical objective that first *minimizes* the number of selected patterns, then the number of degenerate positions, and finally the total base complexity. Because CP-SAT is an anytime solver, it returns an optimal solution if it completes the search and provides a feasible solution if the time limit expires before optimality is established. In all cases, the returned solution is guaranteed to satisfy all exact-cover constraints, and when the solver reaches optimality, the result is mathematically proven to be minimal under the objective hierarchy. TAMIPAMI limits CP-SAT to 30 seconds, returning a feasible solution if the time limit is reached; however, in practice, the solver typically converges within a few seconds and returns an optimal solution.

### Performance benchmarking

Automated performance benchmarking was done for both the command line application and the browser. An automated script available in the repository ran all 7 libraries, 10 times recording, run time, memory, and processor usage for the “predict” and “process” steps. We benchmarked client run time, memory, and processor usage in a Chrome instance of TAMIPAMI. This was done with a Python script using the Selenium WebDriver browser automation library. The Streamlit web application and the Chrome bowser were on the same machine. All benchmarking was run on a 2023 MacBook Pro M2 with 16GB RAM (Mac14,9). All benchmarking code is in the TAMIPAMI GitHub repository.

### Comparison HT-PAMDA code

We attempted to compare TAMIPAMI analysis results to the analysis scripts published in the original HT-PAMDA papers [5,7]. The original HT-PAMDA paper includes a GitHub repository with example code (https://github.com/kleinstiverlab/HT-PAMDA) and subsampled example data. The full data appears to be listed under NCBI Bioproject PRJNA60571, which differs from the Bioproject number listed in the original references 5 and 7. Sample names, read names, and sequences in that Bioproject match the example data in the repository, but we analyzed only the data in the HT-PAMDA GitHub repository and TAMIPAMI due to the discrepancy. Their spCAS9 control library and their timepoint 3 forward read libraries were analyzed with TAMIPAMI. Our sequencing data could not be run with HT-PAMDA due to the fixed design on the HP-PAMDA scripts, and our lack of multiple timepoints.

## Data, web application, and code availability

The TAMIPAMI web application is freely available at https://tamipami.che.ufl.edu. The code for the command line application, web application and benchmarking results is at https://github.com/USDA-ARS-GBRU/tamipami. This code has also been archived with Zenodo at DOI: 10.5281/zenodo.16914326. The sequencing data in this paper are available under NCBI Bioproject number PRJNA1298332.

## Declaration of generative AI and AI-assisted technologies in the writing process

During the preparation of this work, the authors used Grammarly, Microsoft Copilot, and Google Gemini in order to check grammar and the clarity of passages. In the writing of software, authors used Google Jules, Microsoft Copilot, Google Gemini, and Qodo Gen for code editing and review. After using these tools/services, the authors reviewed and edited the content as needed and take full responsibility for the content of the published article.

## Acknowledgements

This work was supported by USDA-ARS in-house research project 6066-21310-006-000-D and USDA-ARS Non-Assistance Cooperative Agreement 58-6066-2-044 to the University of Florida. We thank Mohammadreza Ahmadimashhadi for his help with experiments.

