## Supplemental Data for "TAMIPAMI: Software and methods for PAM/TAM identification in CRISPR and OMEGA gene editing systems" (from another version of this preprint: version 1, 2026-05-16)

**The PDF file includes:**

Fig. S1
Table S1

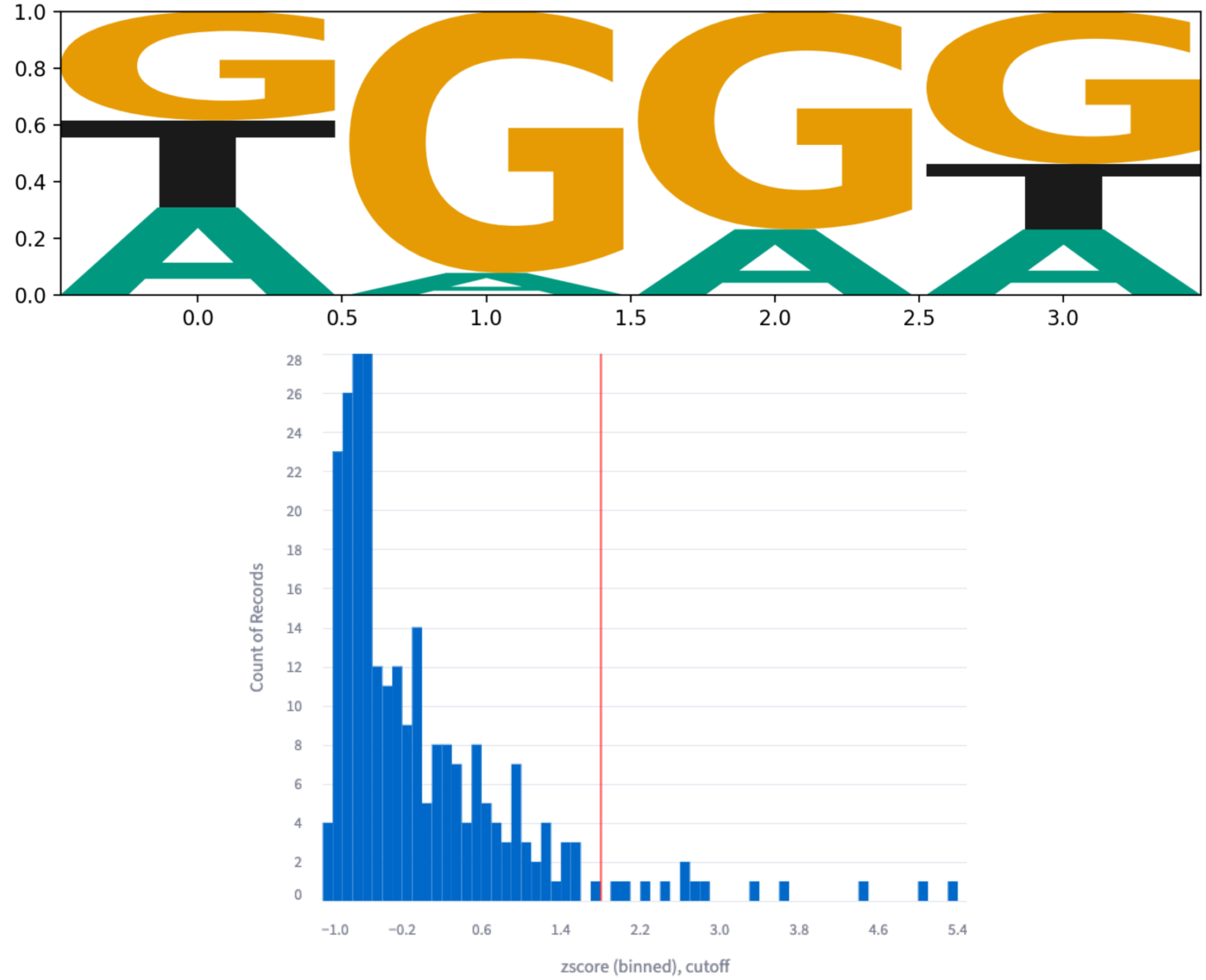

**Figure S1. PAM analysis of SpyCas9 with TAMIPAMI from HT-PAMDA database.** 3-mer motif logos for SpCas9 showing PAM sequences at the 3′ end of the target site and histogram of z score cutoff distribution.

**Table S1.** DNA and RNA sequences used for the study.

| | |
|---|---|
| **crRNA Sequences** | |
| SpCas9 | CGCACCGCTAGCTAATACGACTCACTATAGGGCACGGGCAGCTTGCCGGGTTTTAGAGCTAGAAATAGCAAGTTAAAATAAGGCTAGTCCGTTATCAACTTGAAAAAGTGGCACCGAGTCGGTGC |
| AmaTnpB | TAATACGACTCACTATAGCATCGCTGCGGTGAACATTCGCCGTAGGGCTGAAGTCATGCAGCCGTACGCGGTATAGCGCAGGCTACAACTGCAAGCCCCGCCCTTTAGGGCTGGGGTTCATGACCTGATGGTCCATGTCTGTTA |
| AsCas12a | rUrArArUrUrUrCrUrArCrUrCrUrUrGrUrArGrArUrCrUrGrArUrGrGrUrCrCrArUrGrUrCrUrGrUrUrArCrUrC |
| LbCas12a | rUrArArUrUrUrCrUrArCrUrArArGrUrGrUrArGrArUrCrUrGrArUrGrGrUrCrCrArUrGrUrCrUrGrUrUrArCrUrC |
| Cas12i1 | rArArUrUrUrUrUrGrUrGrCrCrCrArUrCrGrUrUrGrGrCrArCrCrUrGrArUrGrGrUrCrCrArUrGrUrCrUrGrUrUrA |
| BrCas12b | TAATACGACTCACTATAGGAAGGTGGTTAGCTACAGGCTGACCAGTGCAGTTGTGTCATGTGCTACGGTGACCTAACACGTCACTCAGTCACAACGGCTATCTATATTTCCACTAACCAAAGTTAGTGGAAATGTAGATGGTTAGCACCTGATGGTCCATGTCTGTTACTC |
| **Primer Sequences** | |
| Forward Primer | TCGTCGGCAGCGTCAGATGTGTATAAGAGACAGGCATGCCTCGTGACCTGC |
| Reverse Primer | GTCTCGTGGGCTCGGAGATGTGTATAAGAGACAGCGGTATTTCACACCGCATACGTAC |
| **Plasmid Libraries** | |
| Site 2 | Library 160133 / RTW555 |
| Site 4 | Library 160135 / RTW574 |
